# Parallel genomics uncover novel enterococcal-bacteriophage interactions

**DOI:** 10.1101/858506

**Authors:** Anushila Chatterjee, Julia L. E. Willett, Uyen Thy Nguyen, Brendan Monogue, Kelli L. Palmer, Gary M. Dunny, Breck A. Duerkop

**Author notes:** Correspondence: Breck A. Duerkop.

## Abstract

Bacteriophages (phages) have been proposed as alternative therapeutics for the treatment of multidrug resistant bacterial infections. However, there are major gaps in our understanding of the molecular events in bacterial cells that control how bacteria respond to phage predation. Using the model organism *Enterococcus faecalis*, we employed two distinct genomic approaches, transposon (Tn) library screening and RNA sequencing, to investigate the interaction of *E. faecalis* with a virulent phage. We discovered that a transcription factor encoding a LytR family response regulator controls the expression of enterococcal polysaccharide antigen (*epa*) genes that are involved in phage infection and bacterial fitness. In addition, we discovered that DNA mismatch repair mutants rapidly evolve phage adsorption deficiencies, underpinning a molecular basis for *epa* mutation during phage infection. Transcriptomic profiling of phage infected *E. faecalis* revealed broad transcriptional changes influencing viral replication and progeny burst size. We also demonstrate that phage infection alters the expression of bacterial genes associated with intra and inter-bacterial interactions, including genes involved in quorum sensing and polymicrobial competition. Together our results suggest that phage predation has the potential to influence complex microbial behavior and may dictate how bacteria respond to external environmental stimuli. These responses could have collateral effects (positive or negative) on microbial communities such as the host microbiota during phage therapy.

**Importance:** We lack fundamental understanding of how phage infection influences bacterial gene expression and consequently how bacterial responses to phage infection affect the assembly of polymicrobial communities. Using parallel genomic approaches, we have discovered novel transcriptional regulators and metabolic genes that influence phage infection. The integration of whole genome transcriptomic profiling during phage infection has revealed the differential regulation of genes important for group behaviors and polymicrobial interactions. Our work suggests that therapeutic phages could more broadly influence bacterial community composition outside of their intended host targets.

## Introduction

*Enterococcus faecalis* is a member of the healthy human intestinal microbiota (1). *E. faecalis* is also a pathobiont that rapidly outgrows upon antibiotic mediated intestinal dysbiosis to cause disease. *E. faecalis* is associated with nosocomial sepsis, endocarditis, surgical-site, urinary tract and mixed bacterial infections (2, 3). Since the 1980’s, enterococci have been evolving extensive-drug resistance, including resistance to vancomycin and “last-line-of-defense” antibiotics (4–10). In addition, *E. faecalis* can disseminate antibiotic resistance traits to diverse bacteria including other clinically relevant pathogens (11–17). There is an urgent need for new therapeutics that target drug resistant enterococci.

Bacteriophages (phages) are viruses that infect bacteria. Phages are being considered for the treatment of multi-drug resistant (MDR) bacterial infections, including enterococcal infections. Recent studies have demonstrated the potential for phage-based therapies against systemic and biofilm associated enterococcal infections (18–22). The decolonization of intestinal MDR *E. faecalis* may be achieved through the action of phage predation which selects for cell wall variants that are rendered sensitive to antibiotic therapy (23). However, a potential barrier to the wide-spread use of phage therapy against *E. faecalis* is the development of phage resistance. To confront this issue, we must understand the molecular mechanisms used by phages to infect *E. faecalis* and how *E. faecalis* overcomes phage infection to become resistant. Only then can this biology be exploited to better develop phage therapies that mitigate the risk of developing phage resistance.

The study of phage-bacteria interactions has provided key insights into phage infection that could lead to the development of novel antibacterial therapies. Phages replicate in bacteria by hijacking the host cellular machinery to produce phage progeny. To exploit host cell resources, many phages encode auxiliary proteins which are not directly involved in phage genome replication or particle assembly but can modulate bacterial physiology to favor phage propagation (24, 25). The characterization of phage auxiliary proteins may yield tools for curtailing bacterial infections. Additionally, the discovery of phage-modulated host pathways could reveal potential therapeutic targets. Our understanding of global bacterial cellular responses during phage infection is limited to transcriptomic analyses in Gram-negative bacteria, whereas Gram-positive species are understudied (26–31). Therefore, to fill this gap and further define the molecular underpinnings of enterococcal-phage interactions, we have taken a global genomics approach to identify enterococcal factors critical for productive infection by the lytic phage VPE25 (32). To identify bacterial genes essential for VPE25 infection, we screened a low-complexity transposon (Tn) mutant library of *E. faecalis* OG1RF for phage resistance (33). In addition to the known VPE25 receptor (32), transposon sequencing revealed novel *E. faecalis* genes necessary for phage adsorption and optimum intracellular phage DNA replication and transcription. To gain deeper insights into the physiological response of *E. faecalis* during phage infection, we employed temporal transcriptomics of a VPE25 infection cycle. Transcriptomics revealed that VPE25 infection altered the expression of diverse genes involved in protein translation, metabolism, bacterial community sensing, virulence and biofilm formation. Our work indicates that *E. faecalis* reprograms transcription toward stress adaptation in response to phage infection. This suggests that phages may impact the behavior of bacteria in polymicrobial communities including bystanders that are not the intended targets of phage therapy.

## Results

### Transposon sequencing identifies novel genes involved in phage infection of *E. faecalis*

To identify genetic determinants that confer phage resistance in *E. faecalis*, an *E. faecalis* OG1RF transposon library consisting of 6,829 unique mutants was screened by <u>s</u>equence-defined *<u>mar</u>iner* technology <u>tr</u>ansposon <u>seq</u>uencing (SMarT TnSeq) (33). 10^7^ CFU of logarithmically growing *E. faecalis* TnSeq library pool was plated on solid media in the absence and presence of phage VPE25 at a multiplicity of infection (MOI) of 0.1. Cells from the input library prior to plating and cells plated on plates containing no phage were used as controls. Tn insertions in *E. faecalis* genomic DNA were sequenced as described by Dale *et al.* (33). Sequencing reads were mapped to the *E. faecalis* OG1RF genome to identify bacterial mutants with altered phage sensitivity. The relative abundance of 22 *E. faecalis* mutants was enriched (adjusted *P* value < 0.05, log_2_ fold change > 0) in the presence of VPE25 relative to cultures that lacked phage and the input library (Table S1, Fig. 1A). Five of the 22 phage-resistant enriched mutants harbored Tn insertions in OG1RF_10588 (Table S1, Fig. 1A), previously identified to encode the VPE25 receptor <u>P</u>hage Infection <u>P</u>rotein of *E. faecalis* (Pip_EF_) (32). This indicates that the Tn mutant library is an appropriate tool for the discovery of genes involved in phage infection of *E. faecalis*. To gain further insight into the genetic factors that influence *E. faecalis* susceptibility to VPE25, we analyzed several Tn mutants, including OG1RF_10820, OG1RF_10951 (*cscK*), OG1RF_12241, and OG1RF_12435, and three enterococcal polysaccharide antigen (Epa) associated genes, OG1RF_11715 (*epaOX*), OG1RF_11714, and OG1RF_11710 encoding two glycosyltransferases and an O-antigen ligase protein, respectively. (Table S1, Fig. 1A).

**Figure 1.**
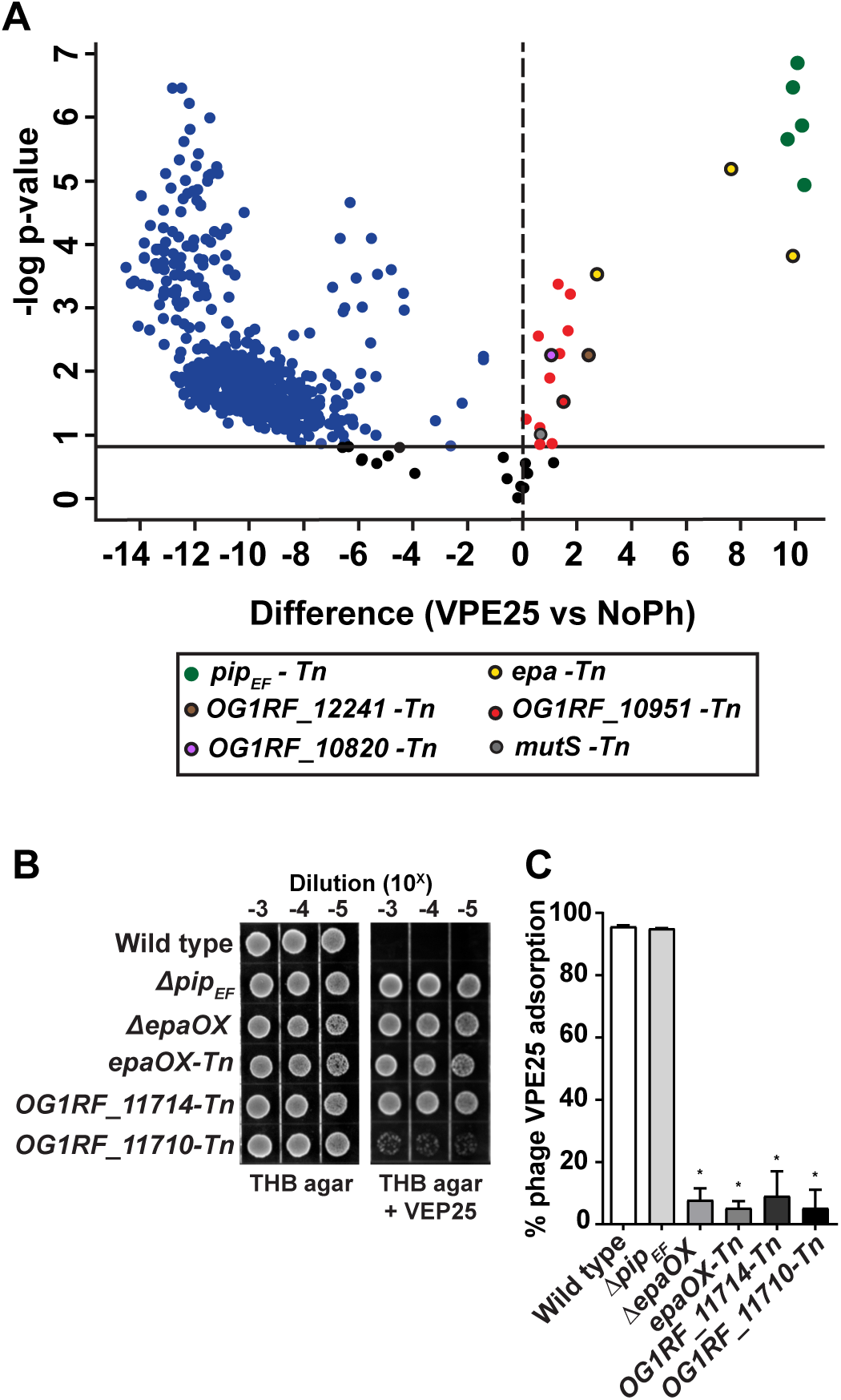
Transposon mutant library screening reveals *E. faecalis* genes important for productive phage infection. **(A)** Volcano plot demonstrating that phage challenge alters the abundance of select mutants from an *E. faecalis* OG1RF Tn library pool compared phage VPE25 naïve *E. faecalis* controls (false discovery rate = 0.05). Phage resistant/tolerant mutants of interest that are enriched upon phage exposure are highlighted: *pip_EF_*-Tn (green), *epa*-Tn (yellow with black outline), *mutS*-Tn (grey with black outline), OG1RF_10820-Tn (purple with black outline), OG1RF_10951-Tn (red with black outline), OG1RF_12241-Tn (brown with black outline). **(B)** Phage VPE25 resistance phenotypes of an isogenic *epa* deletion strain or *epa*-specific Tn mutants serially diluted onto THB agar plates with or without 5 × 10^6^ PFU/ml of VPE25. **(C)** VPE25 efficiently adsorbs to wild type *E. faecalis* OG1RF wild type and an isogenic *Δpip_EF_* deletion strain but not to the various *epa* mutants.

*epa* genes encode proteins involved in the formation of a cell surface-associated rhamno-polysaccharide (34). We and others have previously demonstrated that enterococcal phages unrelated to VPE25 utilize Epa to adsorb and infect *E. faecalis* (23, 35–37). Initial work from our group showed that mutation of the VPE25 receptor Pip_EF_ prevented VPE25 DNA entry into *E. faecalis* V583, yet phages could still adsorb to receptor mutants (32), suggesting that the factors that promote phage infection via surface adsorption remained to be identified. Here, we show that either in-frame deletion of OG1RF_11715 (*epaOX*) (38) or Tn insertions in *epaOX*, OG1RF_11714, and OG1RF_11710 confer phage resistance to VPE25, similar to the *pip_EF_* receptor mutant (Fig. 1B). To assess the role of Epa during VPE25 infection, we investigated the ability of VPE25 to adsorb to wild type *E. faecalis*, the *epaOX* deletion mutant or the three *epa* Tn insertion mutants. Both wild type and the *pip_EF_* mutant strain adsorbed significantly higher amounts of VPE25 as compared to the *epa* mutants (Fig. 1C). Together, these data indicate that *epa*-derived cell wall modifications contribute to VPE25 infection by promoting surface adsorption. This is consistent with previous observations in other lactic acid bacteria that phage infection is a two-step process: first phage must reversibly bind to a cell wall polysaccharide followed by the committed initiation of DNA ejection into the cell (39–41).

In addition to *epa* genes, several Tn mutants whose roles during phage infection were unknown were enriched on VPE25-containing agar compared to uninfected controls (Table S1, Fig.1A). This included OG1RF_10820-Tn, *cscK*-Tn, and OG1RF_12241-Tn. OG1RF_10820 encodes a putative LytR response regulator. Homologs of this protein control multiple cellular processes, including virulence, extracellular polysaccharide biosynthesis, quorum sensing, competence and bacteriocin production (42). OG1RF_12241 is a homolog of the *hypR* (EF2958) gene of *E. faecalis* strain JH2-2 and encodes a LysR family transcriptional regulator. HypR regulates oxidative stress through the *ahpCF* (alkyl hydroperoxide reductase) operon conferring increased survival in mouse peritoneal macrophages (43, 44). Lastly, CscK is a fructose kinase that converts fructose to fructose-6-phosphate for entry into glycolysis (45). Considering that none of these genes had previously been shown to be associated with phage infection, we asked how Tn disruption of these genes influenced phage sensitivity using a time course phage infection assay. In the presence of phage, the optical density of the Tn mutants OG1RF_10820-Tn, *cscK-* Tn, and OG1RF_12241-Tn was maintained constant over time, whereas the growth of wild type and the *pip_EF_* receptor mutant declined or increased over the course of infection respectively (Fig. 2A). Complementation of the Tn mutants with wild type alleles restored phage susceptibility without altering their growth in the absence of phage (Fig. S1A and S1B). To further investigate this phage tolerance phenotype, we asked whether these mutants harbored a defect in phage production. Assessment of the number of phage particles produced during infection showed that OG1RF_10820-Tn, cscK-Tn, and OG1RF_12241-Tn strains produced 10 – fold lower phage particles relative to the wild type strain (Fig. 2B), indicating that the Tn mutants have a defect in phage burst size. In the absence of phage, wild type, *Δpip_EF_*, OG1RF_10820-Tn, *cscK*-Tn, and OG1RF_12241-Tn showed similar growth kinetics (Fig. 2C), suggesting that these mutations do not impair growth in laboratory media. These data indicate that deficiencies in transcriptional signaling and metabolism have a strong impact on lytic phage production in *E. faecalis*.

**Figure 2.**
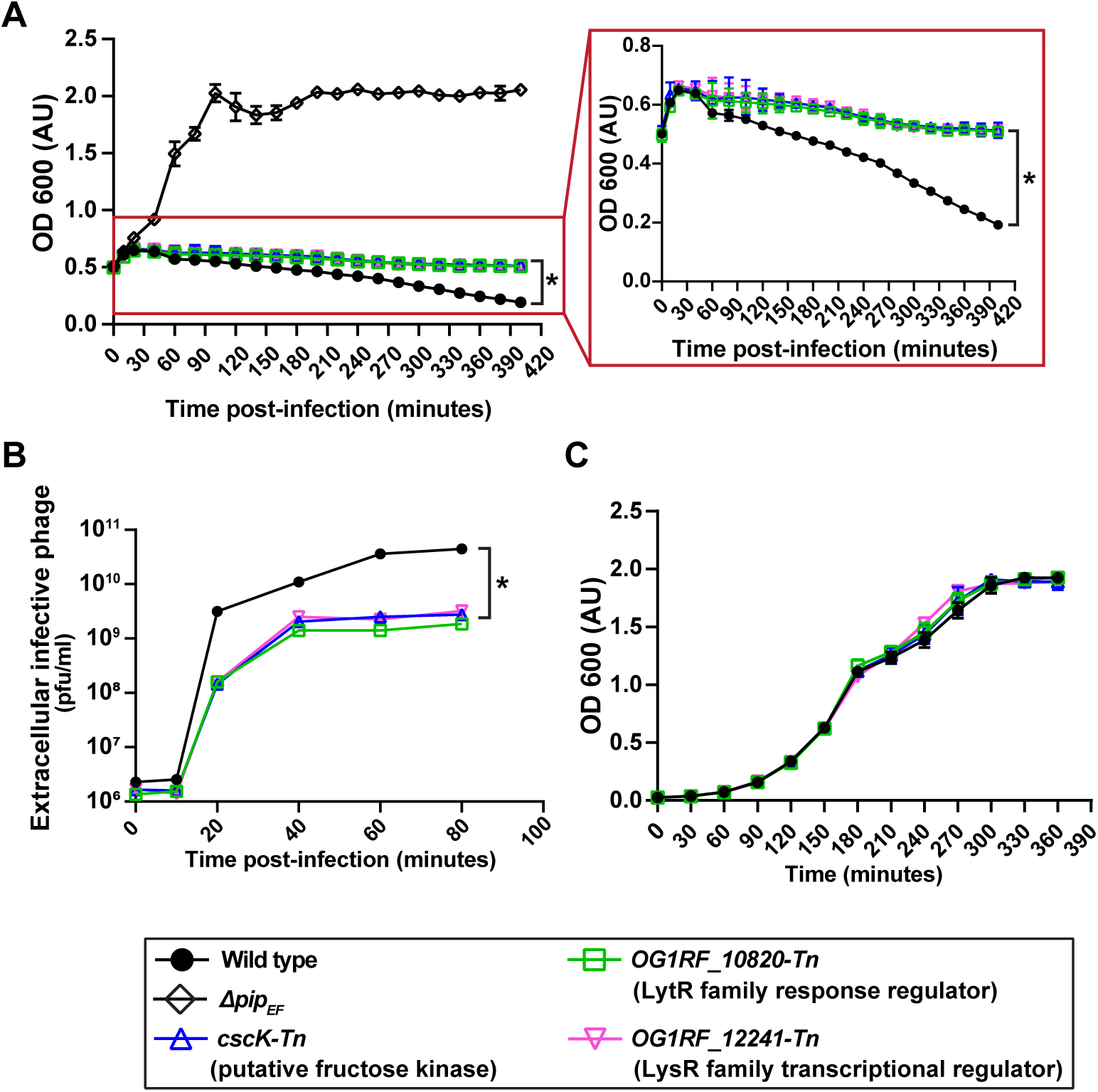
VPE25 mediated killing is halted and phage production is reduced during infection of OG1RF_10820-Tn, cscK-Tn and OG1RF_12241-Tn transposon mutants. **(A)** VPE25 killing curves and **(B)** VPE25 particle production kinetics using the indicated *E. faecalis* transposon mutant strains compared to the wild type or *Δpip_EF_* deletion strains. The inset highlights the delayed lysis phenotype of the transposon mutants relative to wild type. **(C)** Growth curves of all the strains in the absence of VPE25. Data show three independent experiments combined and presented as the mean with standard deviation. \**P* < 0.0001 by two-way analysis of variance (ANOVA).

Since we observed that OG1RF_10820-Tn, cscK-Tn, and OG1RF_12241-Tn mutants had a reduced phage burst, releasing fewer viral particles relative to the wild type, we queried the status of viral transcription and replication in these mutants. Whole genome transcriptomic analysis of phage infected *E. faecalis* OG1RF cells indicated that the VPE25 genes *orf76*, *orf111* and *orf106* are highly expressed throughout the course of phage infection (Table S2A). However, the transcripts of these genes were less abundant in the Tn mutants OG1RF_10820-Tn, *cscK*-Tn, and OG1RF_12241-Tn (Fig. S2A). Additionally, phage DNA replication is delayed in these Tn insertion mutants as judged by significantly lower copy number of phage DNA relative to the wild type strain (Fig. S2B). These data indicate that OG1RF_10820-Tn, *cscK*-Tn, and OG1RF_12241-Tn mutants restrict VPE25 DNA replication which diminishes phage particle production.

### Mutation of *OG1RF_10820* alters *epa* variable gene expression negatively impacting phage infection

To further assess the roles of the *OG1RF_10820*, *cscK*, and *OG1RF_12241* genes during VPE25 infection, we performed phage adsorption assays with these strains. OG1RF_10820-Tn, which harbors a Tn disrupted *lytR* response regulator gene, adsorbed 40% less phage compared to the wild type control (Fig. 3A). Of the three Tn mutants, this was the only mutant that adsorbed less phage compared to the wild type control. LytR type response regulators have been implicated in the biosynthesis of extracellular polysaccharides, including alginate biosynthesis in *Pseudomonas aeruginosa* (42, 46, 47). Since phage adsorption of *E. faecalis* is facilitated by Epa, we measured *epa* gene expression in the OG1RF_10820-Tn background. The *epa* locus consists of core genes (*epaA* – *epaR*) that are conserved in *E. faecalis*, followed by a group of strain-specific variable genes that reside downstream of the core genes (37, 48). The expression of *epa* variable genes *epaOX*, *OG1RF_11714*, and *OG1RF_11710* were reduced in the absence of OG1RF_10820 (*lytR*) during logarithmic and stationary phase growth (Fig. 3B). In contrast, OG1RF_10820 disruption did not alter the expression of core *epa* genes (Fig. S3). Collectively, these results indicate that mutation of the *lytR* homolog hinders optimum binding of VPE25 by downregulating *epa* variable genes thereby modifying the polysaccharide decoration of the core Epa structure (49).

**Figure 3.**
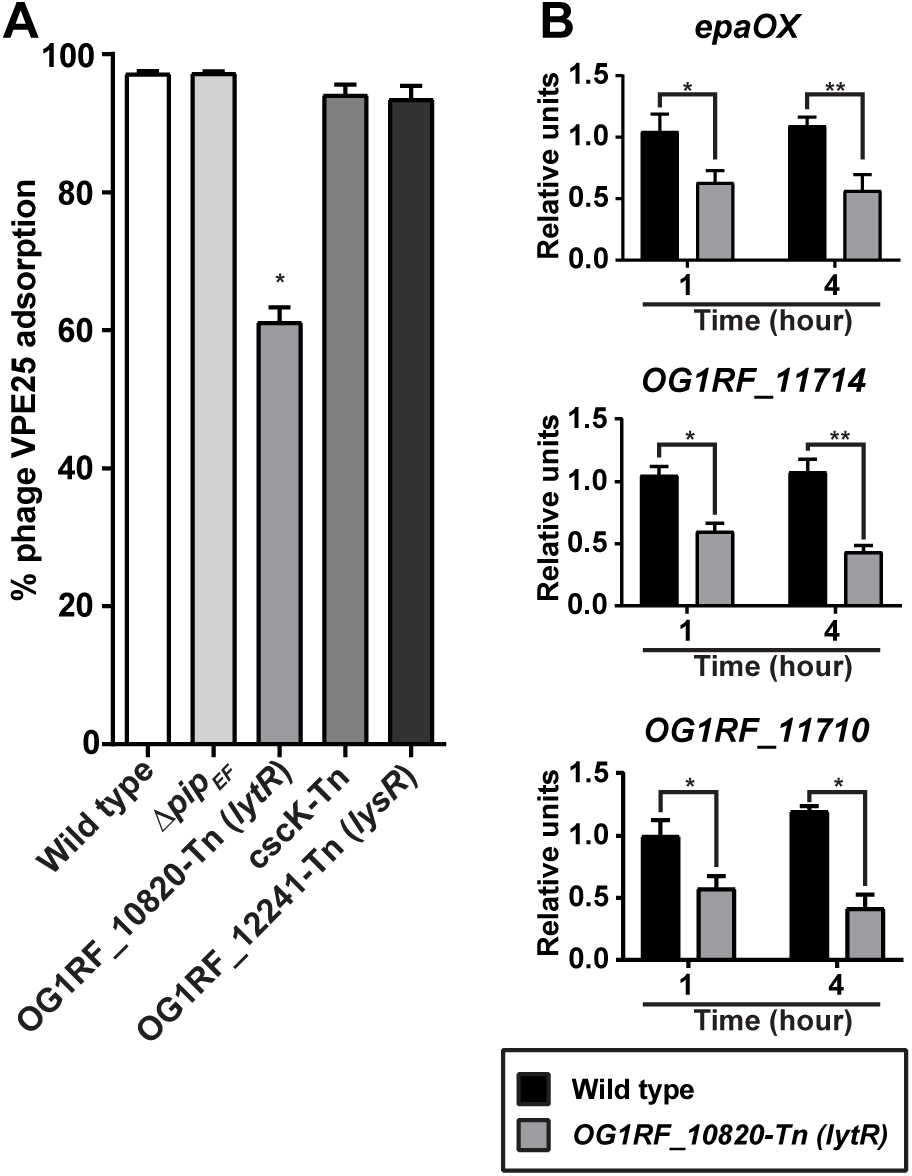
Mutation of a *lytR* homolog downregulates the expression of *epa* variable genes leading to decreased VPE25 adsorption. **(A)** Phage adsorption assay showing that the OG1RF_10820-Tn mutant strain is defective for VPE25 attachment relative to the wild type strain. Disruption of *OG1RF_10820* (*lytR*) leads to reduced expression of three *epa* variable genes **(B)** *epaOX* (upper), *OG1RF_11714* (middle) and *OG1RF_11710* (lower). The data are represented as the fold change of normalized mRNA in comparison to wild type during both logarithmic (1 hr) and stationary phase (4 hr) growth. The data show the average of three biological replicates ± the standard deviation. \**P* < 0.01, \*\**P* < 0.001 by unpaired Student’s t-test.

### Hypermutator strains defective in mismatch repair facilitate acquisition of phage resistance in *E. faecalis*

Transposon mutant OG1RF_12435-Tn, with an insertion in the DNA mismatch repair (MMR) gene *mutS,* was significantly overrepresented during VPE25 infection (Table S1, Fig. 1A). The MMR genes *mutS* and *mutL* correct replication associated mismatch DNA base errors (50). We discovered that VPE25-mediated lysis of the *mutS*-Tn-E (OG1RF_12435-Tn carrying the empty plasmid pAT28) and *mutL*-Tn-E (OG1RF_12434-Tn carrying the empty plasmid pAT28) mutants closely resembled wild-type lysis kinetics for ∼4 hours post-infection and released similar numbers of phage particles (Fig. 4A and 4B). However, these mutator strains eventually started to recover and escape infection suggesting that the mutator phenotype gives rise to phage resistance (Fig. 4A). Introduction of the wild type *mutS* and *mutL* genes cloned into plasmid pAT28 (*mutS*-Tn-C and *mutL*-Tn-C) restored the wild type phage susceptibility phenotype (Fig. 4A). In the absence of phage, the wild type, *mutL*-Tn and *mutS*-Tn strains grew similarly, suggesting that hypermutator strains do not harbor growth defects *in vitro* (Fig. 4C).

**Figure 4.**
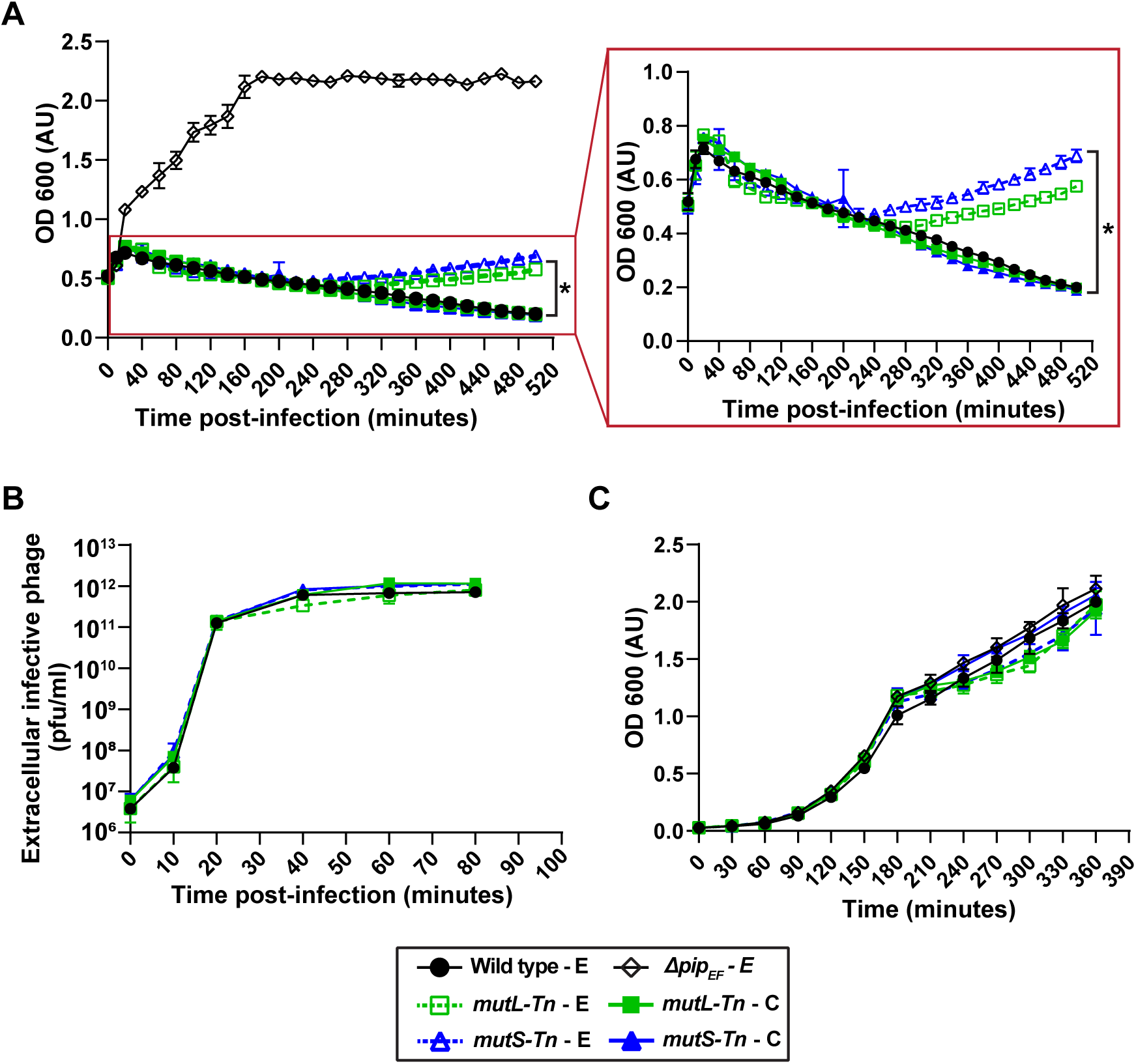
Emergence of phage resistance in *E. faecalis mutL-Tn* and *mutS-Tn* strain backgrounds. **(A)** Culture density of *E. faecalis mutL*-Tn-E and *mutS*-Tn-E strains declined similar to the wild type-E strain following VPE25 infection. However, VPE25 resistance gradually emerged in the *mutL*-Tn-E and *mutS*-Tn-E strain backgrounds as indicated by an increase in cell density following VPE25 infeciton. **(B)** *mutL*-Tn-E, *mutS*-Tn-E and wild type-E strains release equivalent number of phages (PFU/ml) during the course of infection, and **(C)** grow similarly in the absence of phage. Data are show the average of three biological replicates ± the standard deviation. (E, empty vector; C, complemented). \**P* < 0.0001 by two-way analysis of variance (ANOVA).

To confirm that the *mutS*-Tn and *mutL*-Tn strains accumulate phage resistant isolates during phage exposure, we performed phage infection assays using colonies of *mutS*-Tn and *mutL*-Tn grown overnight on agar plates in the absence and presence of VPE25. Consistent with our previous data, all *mutS*-Tn and *mutL*-Tn mutant colonies selected from agar plates lacking phage were initially phage sensitive, and over time became phage resistant (Fig. S4A and S4C). In contrast, the *mutL-*Tn and *mutS-*Tn colonies acquired from the phage containing plates were phage resistant and had similar growth kinetics to the *pip_EF_* receptor mutant in the presence of VPE25 (Fig. S4A and S4C). All isolates grew similarly in the absence of phage (Fig. S4B and S4D). To gain insight into the basis of acquired phage resistance in the mismatch repair mutant backgrounds we compared the phage adsorption profiles of the different strains. The *mutL*-Tn and *mutS*-Tn mutants that were not pre-exposed to VPE25 adsorbed phage at 70 – 80% efficiency, whereas *mutL-*Tn and *mutS-*Tn colonies chosen from phage containing agar plates displayed a severe phage adsorption defect (Fig. S4E and S4F). Our data show that phage treatment *leads to the selection and growth of phage-adsorption deficient isolates from mutL-Tn and mutS-Tn* mutator cultures, most likely through mutations in *epa* variable genes. This also suggests that *epa* variable genes may be a hotspot for mutation in *E. faecalis*.

### VPE25 infection drives global gene expression changes in *E. faecalis*

To study temporal changes in *E. faecalis* gene expression during phage infection, we infected logarithmically growing *E. faecalis* with VPE25 at an MOI of 10. The cell density of infected *E. faecalis* cultures was comparable to the uninfected control cultures during the first 10 min of infection (Fig. 5A). Between 10 and 20 min post infection the VPE25 burst size increased as the cell density of the infected culture declined (Fig. 5A and 5B). VPE25 particle numbers plateaued 30 min post infection, and there was no significant increase in phage output between 30-50 minutes of infection (Fig. 5B).

**Figure 5.**
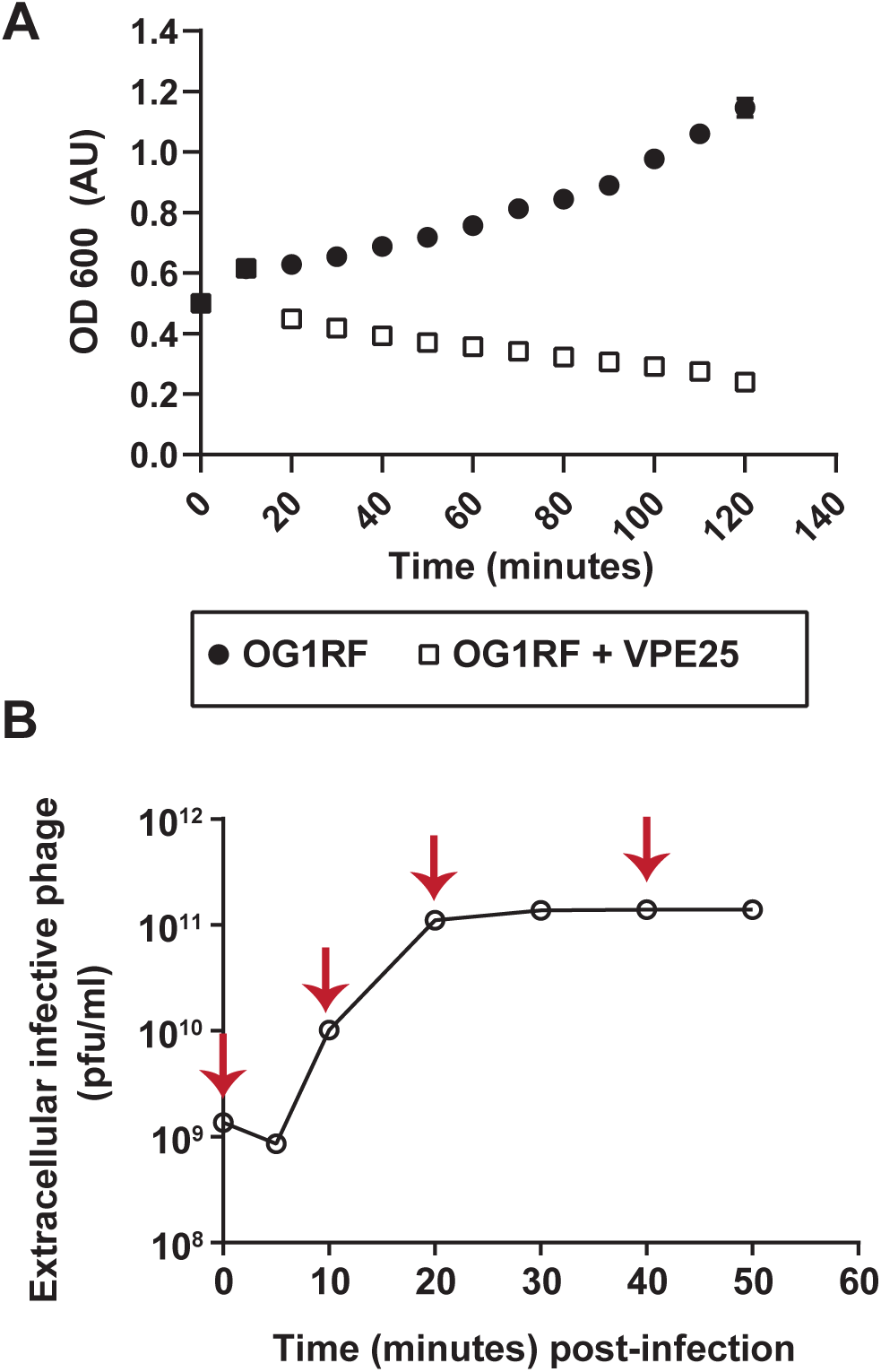
Bacterial growth curve and one – step phage burst kinetics. (A) Optical density of *E. faecalis* OG1RF cultures in the presence and absence of VPE25 infection (MOI = 10). (B) One– step VPE25 growth curve during the infection of *E. faecalis* OG1RF. The red arrows indicate the time points selected for transcriptome analysis. Data from three independent experiments are combined and presented as the mean with standard deviation.

To investigate the transcriptional response of *E. faecalis* during VPE25 infection we collected cells at several time points during distinct phases of the VPE25 infection cycle and performed RNA-Seq. Samples were collected at 10, 20, and 40 minutes post-infection representing the early, middle and late phase of the VPE25 infection cycle, respectively (Fig. 5A and 5B). Hierarchical clustering of differentially expressed *E. faecalis* genes during VPE25 infection compared to uninfected controls revealed unique gene expression patterns at each time point (Fig. 6A and 6B, Table S2B). Gene expression patterns grouped into three distinct clusters that correlated with the early, middle, and late stages of phage infection (Fig. 6A and 6B). Phages rely on host cell resources for the generation of viral progeny. GO and KEGG enrichment analysis showed that phage infection influenced numerous *E. faecalis* metabolic pathways, including amino acid, carbohydrate, and nucleic acid metabolism (Table S2B).

**Figure 6.**
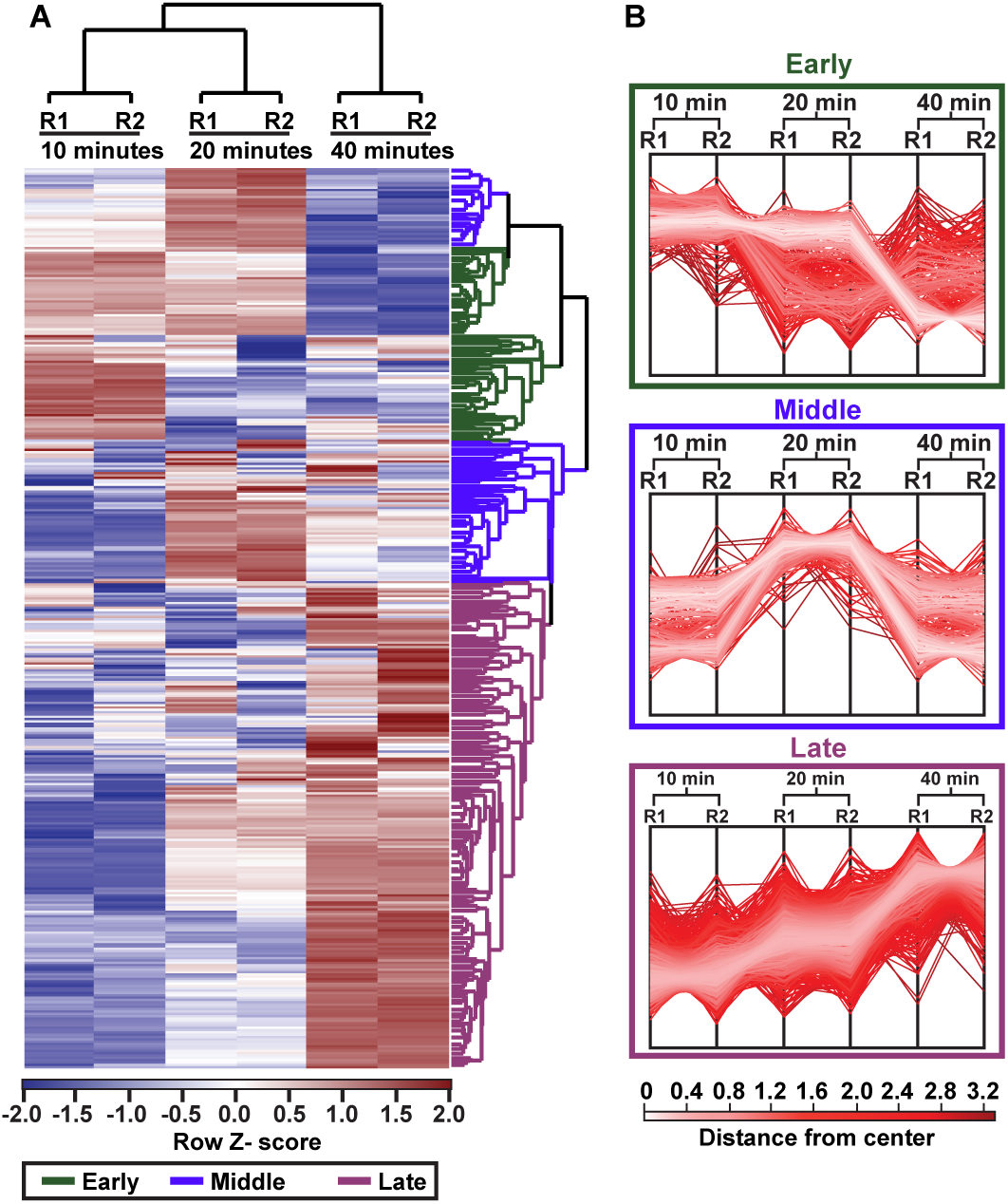
Global transcriptomic profile of *E. faecalis* OG1RF in response to VPE25. **(A)** Hierarchical clustering of differentially expressed *E. faecalis* transcripts at 10, 20 and 40 minutes post VPE25 infection in comparison to uninfected controls from each time point. R1 and R2 designate two independent biological replicates. The transcripts broadly cluster into early (green), middle (blue), and late (magenta) expressed genes. **(B)** The profile plots of the early (top panel), middle (central panel) and late (bottom panel) clusters are shown. Each line indicates a gene within a cluster and the color intensity is calculated based on the distance from the center value in that cluster.

Approximately 54% of the *E. faecalis* genome was differentially expressed (*P* < 0.05) relative to uninfected controls, with 692 downregulated and 731 upregulated genes over the course of phage infection (Fig. S5A and S5C). 37 of the 692 downregulated genes were repressed throughout the course of phage infection and are broadly categorized as ribosome biogenesis and bacterial translation genes (Fig.S5B, Table S3A), indicating that VPE25 modulates host protein biogenesis to prevent bacterial growth and promote viral replication. In contrast, expression of 110 genes belonging to DNA repair pathways, amino acyl-tRNA biosynthesis and carbohydrate metabolism were significantly upregulated throughout the phage infection cycle (Fig. 5D, Table S3B). The induction of DNA stress response genes suggests that *E. faecalis* cells activate DNA defense mechanisms to counteract phage driven DNA damage.

Next, we assessed the transcriptome of VPE25 during infection. We detected 132 differentially expressed VPE25 transcripts by comparing the average read counts of individual genes at 10, 20 and 40 min relative to the start of infection (0 min). Hierarchical clustering grouped differentially expressed genes into early and late genes based on their distinct temporal expression patterns. The transcripts of 78 early genes, including those predicted to be involved in nucleotide biosynthesis and replication, accumulated during the first 10 min of infection (Fig. S6A, Table S2A). In contrast, late genes encoding phage structural components, DNA packaging and host cell lysis were induced by 40 min of infection (Fig. S6A, Table S2A). Approximately 90 genes (68% of the VPE25 genome) annotated as hypothetical in the VPE25 genome were expressed during the early or late phase of infection (Fig. S6A, Table S2A), indicating that the majority of actively transcribed genes during VPE25 infection have no known function.

### Phage infection modulates *E. faecalis* genes involved in group interactions

Our transcriptomic data indicate that VPE25 infection causes a shift in the expression pattern of genes involved in pathways unrelated or peripheral to host metabolism and macromolecule biosynthesis (Fig. S5B and S5D). Most notably, we observed that phage infection led to a significant reduction in the expression of *fsr* quorum sensing genes and in the induction of type VII secretion system genes (T7SS) (Fig. S6B).

The *E. faecalis fsr* quorum sensing system is critical for virulence and biofilm formation in different animal models, including mouse models of endocarditis and peritonitis (51–55). The *fsr* quorum sensing system is comprised of the *fsrA*, *fsrBD* and *fsrC* genes (56–58). *fsrA* encodes a response regulator that is constitutively expressed. *fsrBD* and *fsrC* encode the accessory protein, pheromone peptide and the membrane histidine kinase required for functional quorum sensing regulated gene expression in *E. faecalis*. Quantitative real-time PCR (qPCR) confirmed that *fsrBD* and *fsrC* are repressed throughout VPE25 infection, with the strength of repression increasing over time (Fig. 7A). There was negligible impact on *fsrA* mRNA levels (Fig. 7A) consistent with its behavior as a constitutively expressed gene. qPCR analysis revealed several Fsr-controlled genes to be differentially expressed during phage infection. Fsr-dependent virulence factors, including *gelE*, *sprE*, *OG1RF_10875* (*EF1097)*, and *OG1RF_10876* (*EF1097b*) genes (59) were all significantly downregulated during phage infection relative to uninfected controls (Fig. 7A-B). The *fsr* regulon is also an activator and repressor of several metabolic pathways (60). Our data demonstrates that the levels of Fsr-activated genes involved in the phosphotransferase sugar transport system (OG1RF_10296 and OG1RF_10297) are reduced, whereas negatively regulated genes in the *fsr* regulon, such as *eutB*, *eutC* and *eutH* genes involved in ethanolamine utilization are derepressed during VPE25 infection (Fig. 7B and 7C). These data suggest that VPE25 attenuates the FsrABDC system, consequently impacting *E. faecalis* virulence and metabolism.

**Figure 7.**
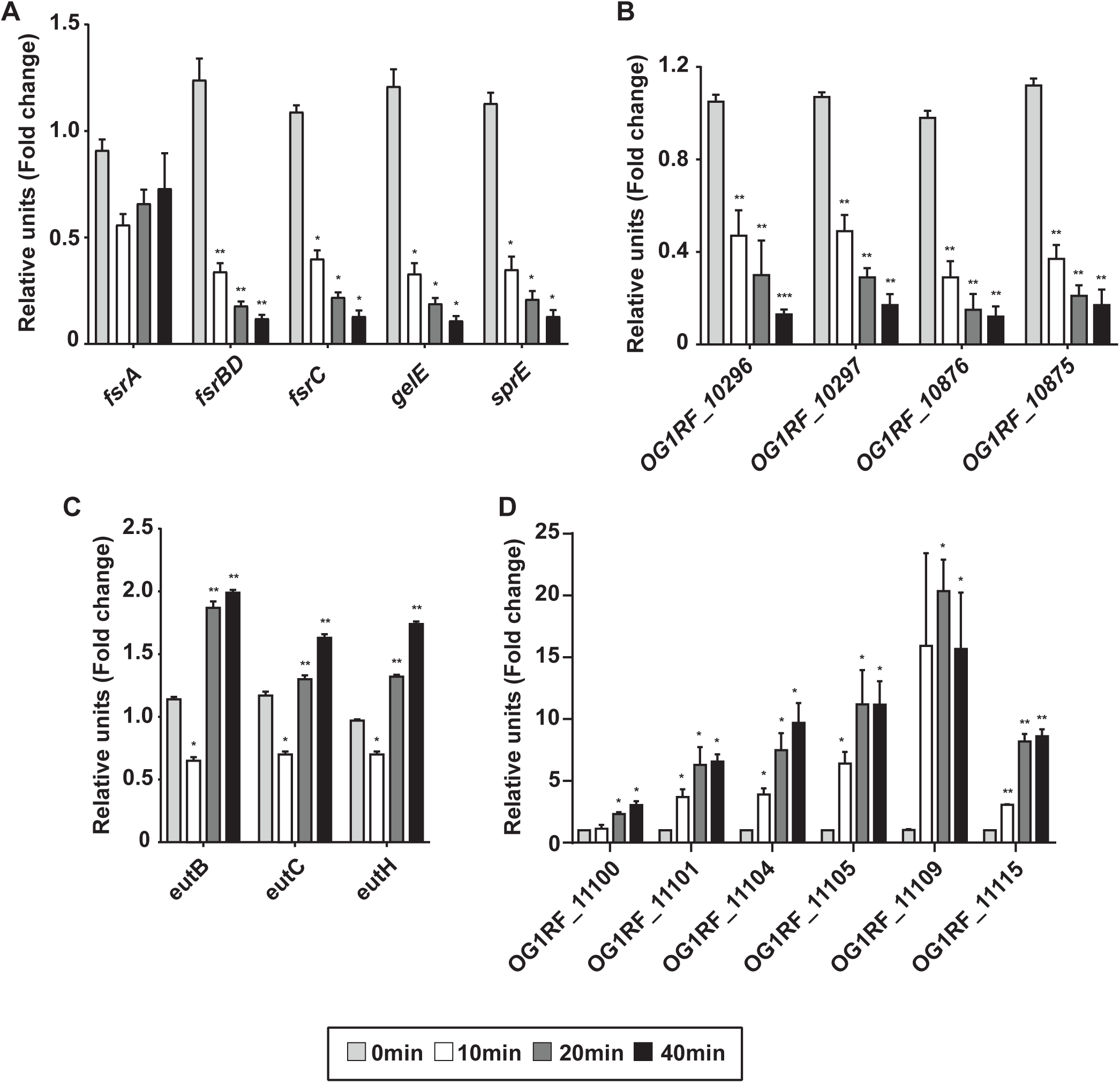
Quantitative PCR confirms altered expression of bacterial quorum sensing and T7SS genes during VPE25 infection. mRNA transcript levels of quorum sensing regulon genes, including **(A)** *fsr* regulatory genes, **(B)** *fsr* – induced genes, and **(C)** *fsr* – repressed genes are differentially expressed during VPE25 infection. **(D)** Progression of the lytic cycle induces the expression of T7SS genes. Expression is the fold change relative to untreated samples at the same time points. Data represent the average of three replicates ± the standard deviation. \**P* < 0.01 to 0.0001 and \*\**P* < 0.00001 by unpaired Student’s t-test.

Divergent forms of the T7SS (also known as the Ess/Esx system) are widely distributed in Gram-positive bacteria and the T7SS has been extensively studied in the Actinobacterium, *Mycobacterium tuberculosis* (61–66). To date, enterococcal T7SS loci remain uncharacterized. Consistent with our transcriptomic data (Fig. S6B), we observed that phage infection induces genes in the *E. faecalis* T7SS locus, including *OG1RF_11100* (*esxA*), *OG1RF_11101* (*essA*), *OG1RF_11104* (*essB*), *OG1RF_11105* (*essC1*), *OG1RF_11109* and *OG1RF_11115* (*essC2*) (Fig. 7D). The *esxA* and *OG1RF_11109* genes encode potential WXG100 domain containing effector and LXG-domain toxin proteins, respectively. The secretion of T7SS factors is dependent on the EssB transmembrane protein, and FtsK/SpoIIIE ATPases encoded by the *essC1* and *essC2* genes. The T7SS in Gram-positive bacteria is involved in immune system activation, apoptosis of mammalian cells, bacterial cell development and lysis, DNA transfer and bacterial interspecies interactions (62, 67–77).

To investigate whether phage-mediated expression of *fsrABDC* and the T7SS genes are VPE25 specific, we examined the expression patterns of a subset of these genes in *E. faecalis* when infected with the unrelated phage NPV-1. mRNA levels of *fsrB* and *sprE* were reduced whereas expression of the T7SS genes were elevated during NPV-1 infection (Fig. S6C), suggesting that phage specific control of quorum sensing and T7SS expression is not restricted to VPE25 infection. Together these findings indicate that phage infection of *E. faecalis* has the potential to influence bacterial adaptation in polymicrobial communities or during mixed bacterial species infections.

## Discussion

Characterizing bacterial responses to phage infection is important for understanding how phages modulate bacterial physiology and will inform approaches toward effective phage therapies against MDR bacteria. Commonly used approaches to identify phage resistant bacteria often yield information restricted to phage receptors and/or adsorption mechanisms. While useful, these approaches often overlook more subtle interactions that drive the efficiency of phage infection and phage particle biogenesis. Additionally, these approaches provide minimal information on how bacteria sense and respond to phage infection. These are important gaps in knowledge considering the heightened interest in utilizing phages as clinical therapeutics against difficult to treat bacterial infections. To begin to address these knowledge gaps we have studied the model Gram-positive commensal and opportunistic pathogen *E. faecalis* when infected with its cognate lytic phage VPE25. We have discovered several novel bacterial factors that are indispensable for efficient VPE25 infection of *E. faecalis*. In addition, we have uncovered key insights into the molecular events that are triggered in *E. faecalis* cells during phage infection. Importantly, our work shows that *E. faecalis* alters the expression of genes associated with environmental sensing and group interactions during phage infection. Such a response may have unexpected consequences in polymicrobial communities, and our work sets the stage for studying how phage therapies may impact non-target bacteria in the microbiota.

TnSeq identified numerous *E. faecalis* genes that govern VPE25 susceptibility. Mutations in *epa* variable genes conferred VPE25 resistance by preventing phage adsorption, similar to other *E. faecalis* phages (23, 35–37). We discovered that phage infection of mismatch repair gene mutants results in the emergence of phage adsorption deficiencies, thus the mismatch repair system likely fails to correct DNA damage of *epa* genes during phage infection. This suggests that *epa* genes may be a hotspot for mutation. TnSeq also enabled the discovery of the LytR-domain transcription factor encoded by OG1RF_10820 as a regulator of *epa* variable locus gene expression (Fig. 3B). Considering an *epa* mutant strain of *E. faecalis* is defective in colonization and outgrowth during antibiotic selection in the intestine (23), we believe that future investigation of LytR mediated regulation of *epa* and potentially other genomic loci could guide the development of effective therapeutics to control *E. faecalis* colonization and infections.

This work has discovered bacterial metabolic and oxidative stress response genes that are important for phage infection. Decreased phage replication coupled with lower burst size of VPE25 in the fructose kinase mutant (*cscK*) suggests that VPE25 relies on host carbohydrate metabolism to support viral progeny formation. The reliance of VPE25 on host CscK for viral propagation is corroborated by RNA-Seq data showing broad induction of bacterial host carbohydrate metabolism genes during VPE25 infection. Further, we observed that VPE25 lytic growth leads to a gradual rise in the OG1RF_12241 transcripts which encode the LysR homolog HypR, a regulator of oxidative stress. Phage infection enhanced the transcript levels of three other oxidative stress response genes, including OG1RF_10348, OG1RF_10983, and OG1RF_11314 encoding superoxide dismutase (*sodA*), NADH peroxidase (*npr*) and catalase (*katA*), respectively (Table S2B). In *Campylobacter jejuni*, mutations in the LysR regulated gene *ahpC*, as well as *sodB* and *katA* resulted in reduced plaquing efficiency by the phage NCTC 12673 (27). We hypothesize that phage tolerance during hypersensitivity to oxidative stress could be detrimental to *E. faecalis* and targeting such pathways could be used to control *E. faecalis* colonization.

Our data indicate that putative T7SS genes are activated in response to phage infection. Although *E. faecalis* T7SS remains poorly characterized, the *Staphylococcus aureus* T7 system has been demonstrated to defend cells against neutrophil assault, enhance epithelial cell apoptosis and is critical for virulence (68, 69, 78, 79). Additionally, *S. aureus* T7SS maintains membrane homeostasis and is involved in the membrane stress response (69, 80). The finding that *E. faecalis* T7SS genes are induced in response to two different phages suggests phage mediated membrane damage may lead to elevated T7SS gene expression. Finally, the *S. aureus* T7SS nuclease toxin, EsaD, contributes to interspecies competition through growth inhibition of rival strains lacking the EsaG anti-toxin (77). The impact of *S. aureus* T7SS on rival strains and the presence of T7SS genes in environmental isolates (81, 82) suggests a pivotal role of this secretion system in shaping microbial communities. Future studies on *E. faecalis* T7SS will aim to determine how this system influences intra- and/or interspecies competition and niche establishment in polymicrobial environments such as the microbiota.

In contrast to the T7SS, the bacterial population associated quorum sensing *fsr* locus was repressed during VPE25 infection in *E. faecalis*. Gram–negative bacteria can escape phage invasion by quorum sensing mediated downregulation of phage receptor expression or activation of CRISPR-Cas (clustered regularly interspaced short palindromic repeats) immunity (83–86). The *fsr* regulon does not include the receptor (*pip_EF_*), *epa* genes necessary for phage adsorption, or CRISPR-Cas. However, the *fsr* system does contribute to biofilm formation that could potentially deter phage infection (55, 87). VPE25 infection may attenuate *fsr* mediated biofilm formation to favor continued infection of neighboring planktonic cells. On the other hand, phage genomes have been shown to carry enzymes that degrade quorum sensing molecules or anti-CRISPR genes to evade host defense strategies (88, 89). Although such anti-host accessory genes are not evident in the VPE25 genome, it is possible that phage encoded hypothetical genes influence *E. faecalis* quorum sensing and dictate molecular events that favor phage production. Identification of such auxiliary phage proteins could lead to the discovery of potential anti-enterococcal therapeutics.

Integration of global transcriptomics using transposon library screening of VPE25-infected *E. faecali*s has revealed new insights into our understanding of phage-host interactions in enterococci. Together, our results emphasize the importance of *epa* gene regulation, carbohydrate metabolism and the oxidative stress response in successful phage predation. Further, contributions of VPE25 on *E. faecalis fsr* and T7SS genes involved in inter- and intra-bacterial interactions suggests that phage therapy could impact microbial community dynamics in patients undergoing treatment, and such an outcome should be taken into consideration for the development of phage-based therapeutics.

## Materials and methods

### Bacteria and bacteriophages

All bacteria and phages used in this study are listed in Table S4. *E. faecalis* strains were grown with aeration on Todd-Hewitt broth (THB) or THB agar at 37°C. *Escherichia coli* was grown on Lennox L broth (LB) with aeration or on LB agar at 37°C. The following antibiotic concentrations were added to media for selection of *E. coli* or *E. faecalis*: 25 μg/ml fusidic acid, 50 μg/ml rifampin, 750 μg/ml spectinomycin and 20 μg/ml chloramphenicol. Phage sensitivity assays were performed on THB agar supplemented with 10mM MgSO_4_.

### Transposon library screen

10^8^ colony forming units (CFU) of the *E. faecalis* OG1RF pooled transposon library was inoculated into 5 ml of THB and grown with aeration to an optical density of 600 nm (OD_600_) of 0.5. 10^7^ CFU of the library was spread onto a THB agar plate (10 replicates) containing 10mM MgSO_4_ in the absence and presence of 10^6^ plaque forming units (PFU) of VPE25 (MOI = 0.1). After overnight (O/N) incubation at 37°C, bacterial growth from the control and phage containing plates was resuspended in 5 ml of phosphate buffered saline (PBS). Genomic DNA was isolated from the input library and from three biological replicates of phage exposed and unexposed samples using a ZymoBIOMICS™ DNA Miniprep Kit (Zymo Research), following the manufacturers protocol.

### Transposon library sequencing

Library preparation and sequencing was performed by the Microarray and Genomics Core at the University of Colorado Anschutz Medical Campus. A detailed protocol is described by Dale *et al.* (33). Briefly, 100 ng of genomic DNA was sheared to approximately 400 bp fragments and processed through the Illumina TruSeq Nano library enrichment kit. 9 ng of each normalized library was used as PCR template to enrich for the *mariner* transposon junctions using a transposon-specific primer (*mariner*-seq) and the Illumina P7 primer (16 cycles of amplification). The enrichment PCR products were diluted 1:100, and 10 μl was used as template for an indexing PCR of 9 cycles of amplification (TruSeq P5 indexing primer + P7 primer). The final libraries had unique combinations of P5 and P7 indexes suitable for multiplexed sequencing. Sequencing was performed using Illumina NovaSeq 6000 in 150 base paired-end format. Illumina adapter trimming, read mapping to the *E. faecalis* OG1RF reference sequence (NC_017316.1) and statistical analysis of differentially abundant transposon mutants were performed using previously published scripts found here https://github.com/dunnylabumn/Ef_OG1RF_tnseq (33).

### Bacterial spot assay on phage agar plates

O/N bacterial cultures were pelleted and resuspended in SM-plus buffer (100 mM NaCl, 50 mM Tris-HCl, 8 mM MgSO_4_, 5 mM CaCl_2_ [pH 7.4]) and normalized to an OD_600_ of 1.0. 10-fold serial dilutions of the bacterial cultures were spotted onto THB agar plates with or without 5x10^6^ PFU/ml of VPE25. The plates were incubated at 37°C O/N.

### Phage sensitivity and burst kinetic assays

O/N cultures of *E. faecalis* were subcultured to a starting OD_600_ of 0.025 in 25 ml of THB. When the bacterial culture reached mid-logarithmic phase (OD_600_ ∼ 0.5), 10mM MgSO_4_ and VPE25 (MOI of 0.1 or 10) were added. OD_600_ was monitored for ∼ 7 hrs. To investigate if phage progeny were produced and released from the bacterial cells upon VPE25 infection, 250 μl of culture was collected at different time points over the course of infection and thoroughly mixed with 1/3 volume of chloroform. The aqueous phase containing phages was separated from the chloroform by centrifugation at 24,000 × *g* for 1 min and the phage titer was determined using a THB agar overlay plaque assay. Data are presented as the average of three replicates with +/- standard deviation.

### Bacterial growth curves

25 ml of THB was inoculated with O/N cultures of *E. faecalis* to obtain a starting OD_600_ of 0.025. Cultures were incubated at 37° C with aeration. OD_600_ was measured periodically for ∼7 hours. Growth curves are presented as the average of three biological replicates.

### Phage adsorption assay

A bacterial culture grown O/N was pelleted at 3,220 × *g* for 10 min and resuspended to 10^8^ CFU/ml in SM-plus buffer. The cell suspensions were mixed with phages at an MOI of 0.1 and incubated at room temperature without agitation for 10 min. The bacterium-phage suspensions were centrifuged at 24,000 × *g* for 1 min, and the supernatant was collected and phages were enumerated by a plaque assay. SM-plus buffer with phage only (no bacteria) served as a control. Percent adsorption was determined as follows: [(PFU_control_ − PFU_test supernatant_)/PFU_control_] × 100. Data are presented as the average of three replicates +/- standard deviation.

### Complementation of Tn mutants

All PCR reactions used for cloning were performed with high fidelity KOD Hot Start DNA Polymerase (EMD Millipore). Approximately 100 bp of upstream flanking DNA and the coding regions of OG1RF_10820, OG1RF_10951 (*cscK*), OG1RF_12241, OG1RF_12435 (*mutS*) and OG1RF_12434 (*mutL*) were cloned into the shuttle vector pAT28 (90). The primer sequences and restriction enzymes used for cloning are listed in Table S4. Plasmids were introduced into electrocompetent *E. faecalis* cells as previously described (23).

### RNA extraction and quantitative PCR

RNA was extracted from uninfected or VPE25 infected *E. faecalis* by using an RNeasy Mini Kit (Qiagen) with the following modifications. Cell pellets were incubated in 100 µL of 15mg/ml lysozyme (Amersco) for 30 min at room temperature. 700 µl of RLT buffer containing β-mercaptoethanol (manufacturers recommended concentration) was added and the samples were bead beat in Lysing Matix B tubes (MP Bio) at 45 sec intervals for a total time of 4.5 min. Debris was centrifuged at 24,000 × *g* for 1 min and the supernatant was transferred to a fresh tube. 590 µL of 80% ethanol per 760 µL supernatant was added and the entire volume was loaded onto an RNeasy column following the standard Qiagen RNA purification protocol. cDNA was synthesized from 1 µg of total RNA using qScript cDNA SuperMix (QuantaBio) (25°C for 5 minutes, 42°C for 30 minutes and 85°C for 5 minutes). Transcript levels were analyzed by qPCR using PowerUp^TM^ SYBR Green Master Mix (Applied Biosystems) and transcript abundances were normalized to the 16S rRNA transcripts. VPE25 orf_76 copy number was determined by qPCR using orf_76 cloned into pCR4-TOPO™ TA cloning vector (Invitrogen) as a standard. All data are represented as the average of three replicates +/- the standard deviation.

### RNA sequencing and bioinformatics analysis

O/N cultures of *E. faecalis* were subcultured to a starting OD_600_ of 0.025 in 50 ml of THB. When the bacterial culture reached mid-logarithmic phase (OD_600_ ∼ 0.5), 10mM MgSO_4_ and VPE25 (MOI of 10) were added. 4ml of cell suspension was pelleted from the uninfected and infected cultures after 0, 10, 20 and 40 minutes post VPE25 treatment. The pellets were washed with 4ml of PBS three times, followed by a wash with 2ml RNA*later*^TM^ (Invitrogen) and RNA isolation was performed as described above. RNASeq libraries were constructed using the Ribo Depleted library construction kit for Gram-positive Bacteria (Illumina). Sequencing was performed using Illumina NovaSeq 6000 in 150 base paired-end format. RNASeq data were analyzed using Geneious R11. Sequencing reads were mapped to the *E. faecalis* OG1RF (NC_017316.1) and VPE25 (LT546030.1) genomes using Bowtie2. Gene expression values were calculated by reads per kilobase per million which normalizes the raw count by transcript length and sequence depth. Differential expression between two samples was determined using the default Geneious R11 method with median ratios across all the transcripts as the normalization scale. Genes with a fold change of ≥ 2.0 and *P* values ≤ 0.05 were considered significantly differentially expressed. Blast2GO basic tool was used to assign gene ontology (GO) terms to *E. faecalis* OG1RF genes (91). KEGG pathway analysis was performed using the KEGG annotation server (92). The ClueGo plug-in for Cytoscape 3.6.1 (93, 94) was used to visualize gene clustering based on GO terms and KEGG annotations.

### Statistical analysis

Statistical tests were performed using GraphPad – Prism version 8.2.1. For bacterial growth assays in the presence and absence of phage, mutant *E. faecalis* was compared to the wild type using two-way analysis of variance (ANOVA). For qPCR and phage adsorption assays, unpaired Student’s t-tests were used. *P* values are indicated in the figure legends.

### Data availability

The RNA-Seq reads associated with this study have been deposited at the ArrayExpress database at EMBL-EBI under accession number E-MTAB-8546. Tn-Seq reads have been deposited at the European Nucleotide Archive under accession number PRJEB35492.

## Acknowledgments

This work was supported by National Institutes of Health grants R01AI141479 (B.A.D.) and R01AI116610 (K.L.P.). J.L.E.W. was supported by American Heart Association Grant 19POST34450124 / Julia Willett / 2018. We would like to thank Katrina Diener and Monica Ransom from the University of Colorado Anschutz Medical Campus Genomics and Microarray Core for the development of a customized Tn-Seq library preparation protocol.

---

**Figure S1.**
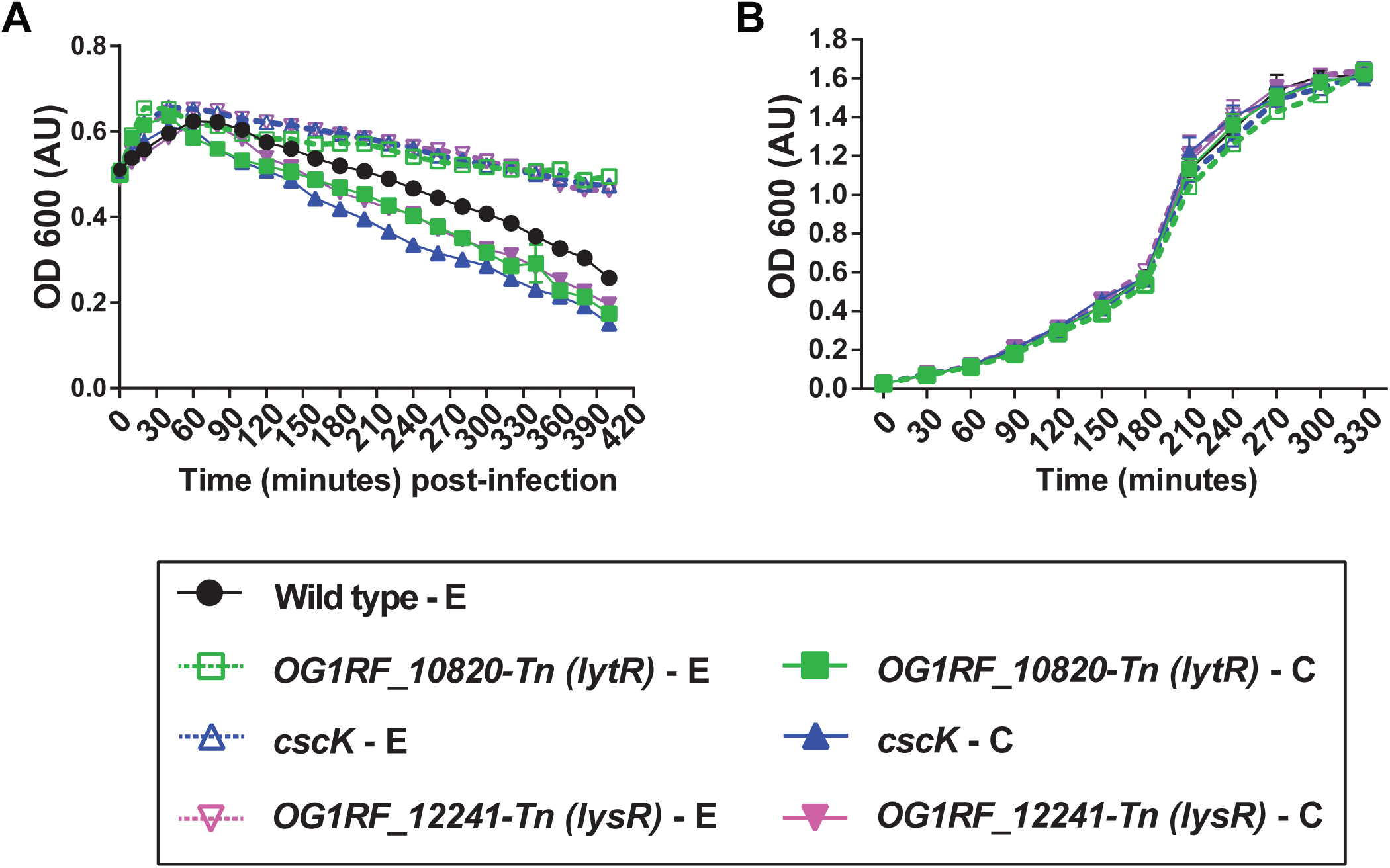
Complementation restores phage sensitivity in OG1RF_10820, *cscK* and OG1RF_12241 Tn mutants. **(A)** Introduction of the wild type allele but not the empty plasmid sensitizes *E. faecalis* Tn mutants to phage VPE25 infection. **(B)** Growth in the absence of VPE25 remains unaltered irrespective of the presence of the empty or complementation plasmid. (E, empty vector; C, complemented).

**Figure S2.**
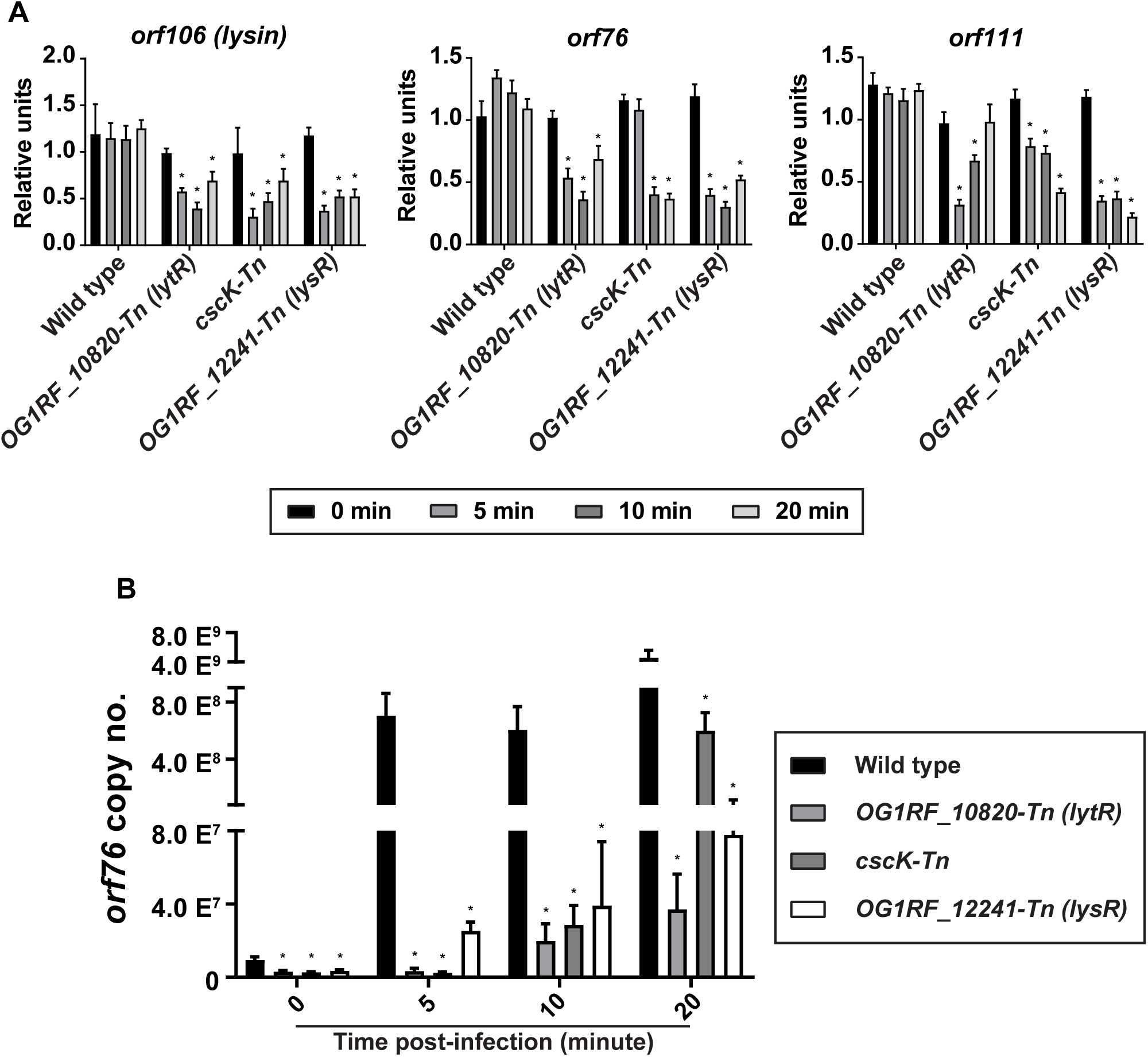
Dampened viral gene expression and DNA replication aids in OG1RF_10820-Tn, *cscK*-Tn and OG1RF_12241-Tn mutant tolerance to VPE25 infection. (A) Effect of various Tn insertion mutations on VPE25 mRNA levels. The data are shown as the fold change of normalized mRNA in comparison to the wild type samples at various time points post-infection. **(B)** VPE25 DNA copy number calculated from an *orf76* standard curve. Data are represented as average of three replicates ± the standard deviation. \**P* < 0.00001 by unpaired Student’s t-test.

**Figure S3.**
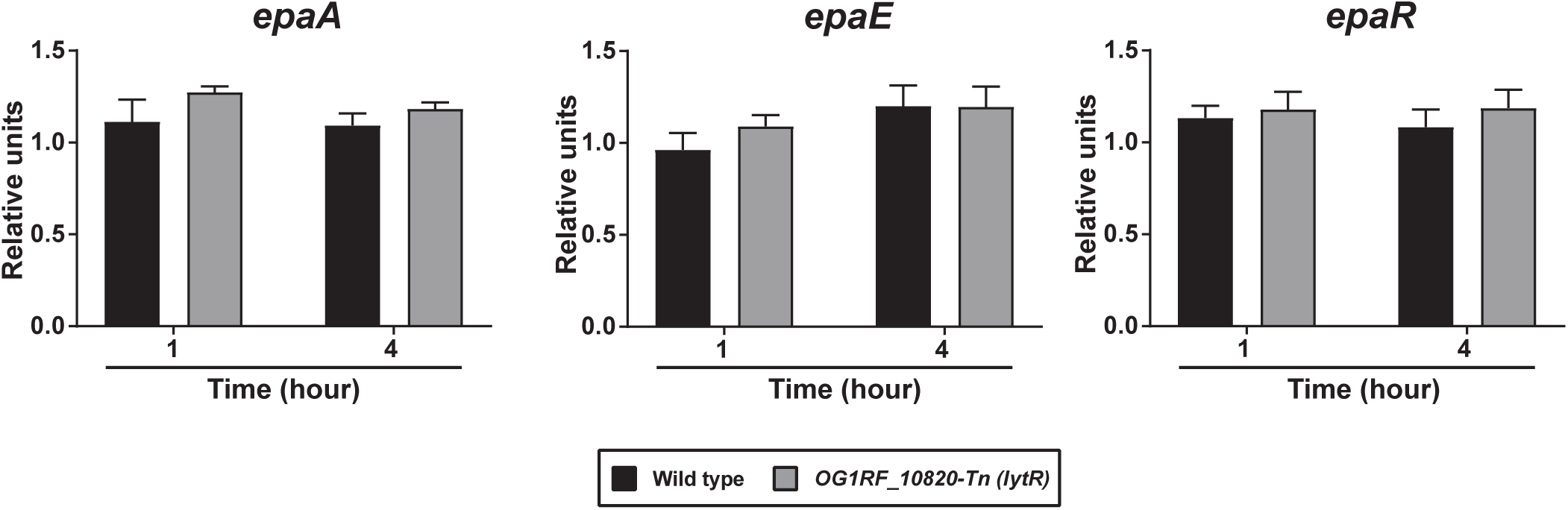
The *lytR* gene does not influence the expression of *epa* core genes. Quantitative PCR demonstrates equivalent mRNA levels of the *epa* genes *epaA*, *epaE*, and *epaR* in wild type and the OG1RF_10820-Tn *E. faecalis* strains. The data are expressed as fold change of normalized mRNA in comparison to the wild type at different time points and represent the average of three biological replicates ± the standard deviation.

**Figure S4.**
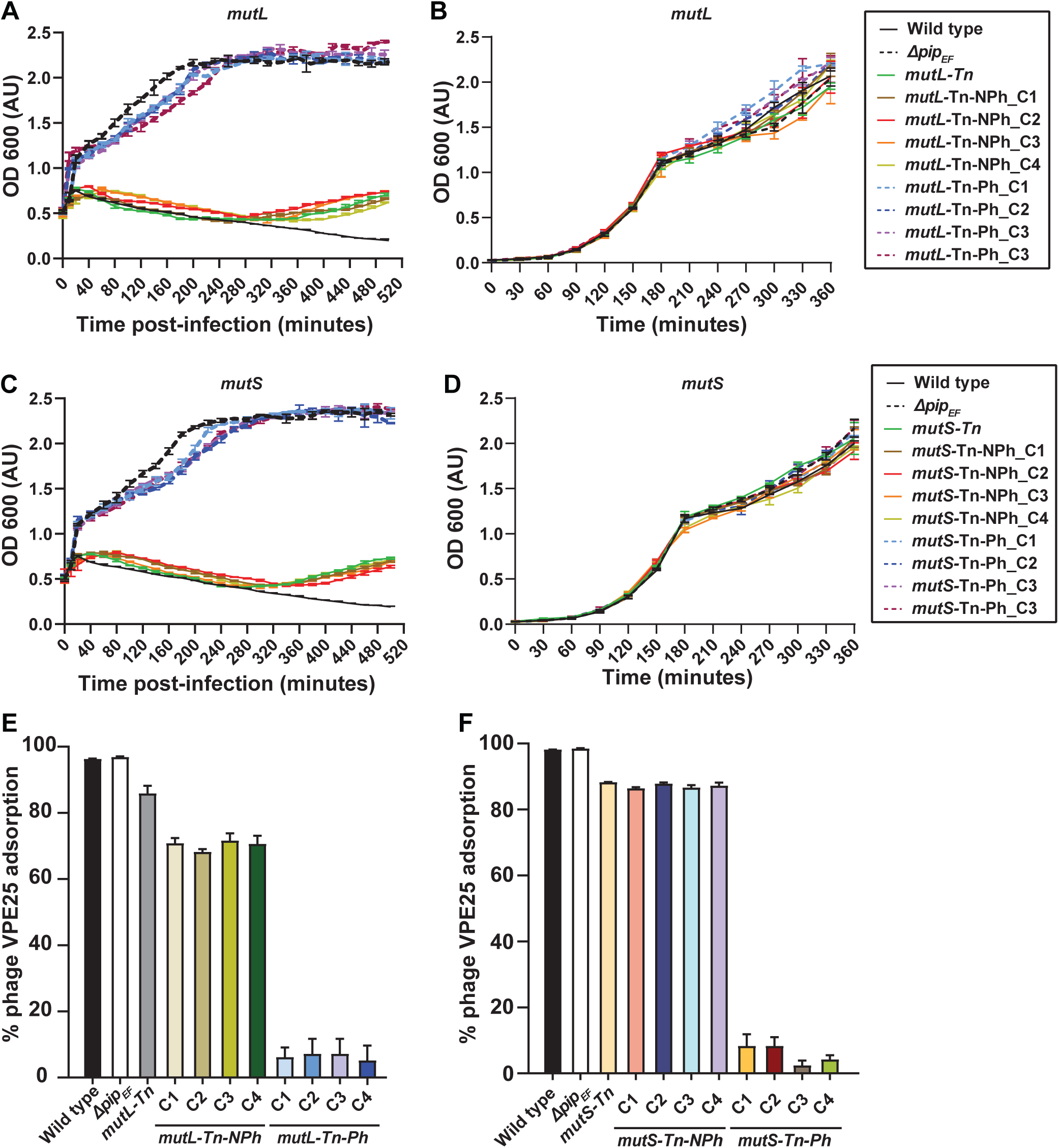
Mutator strains facilitate the acquisition of VPE25 resistance. Growth of wild type, *Δpip*, parent *mutL-Tn*, *mutL-Tn* colonies selected from THB plate without phage (mutL-Tn-NPh_C1 to C4 and mutS-Tn-NPh_C1 to C4) and those selected from VPE25 (5 × 10^6^ PFU/ml) containing THB plates (mutL-Tn-Ph_C1 to C4 and mutS-Tn_C1 to C4) are shown **(A and C)** in the presence and **(B and D)** in the absence of VPE25. The mutL-Tn and mutS-Tn colonies pre-exposed to phage phage behave similar to phage resistant *Δpip*, whereas hypermutator colonies without previous VPE25 challenge gained phage resistance during the course of infection. **(E-F)** All the mutL-Tn-Ph_C1 to C4 and mutS-Tn-Ph_C1 to C4have a defective VPE25 adsorption profile, while colonies selected from no phage plates are able to adsorb ∼ 70-80% of the phages in the assay.

**Figure S5.**
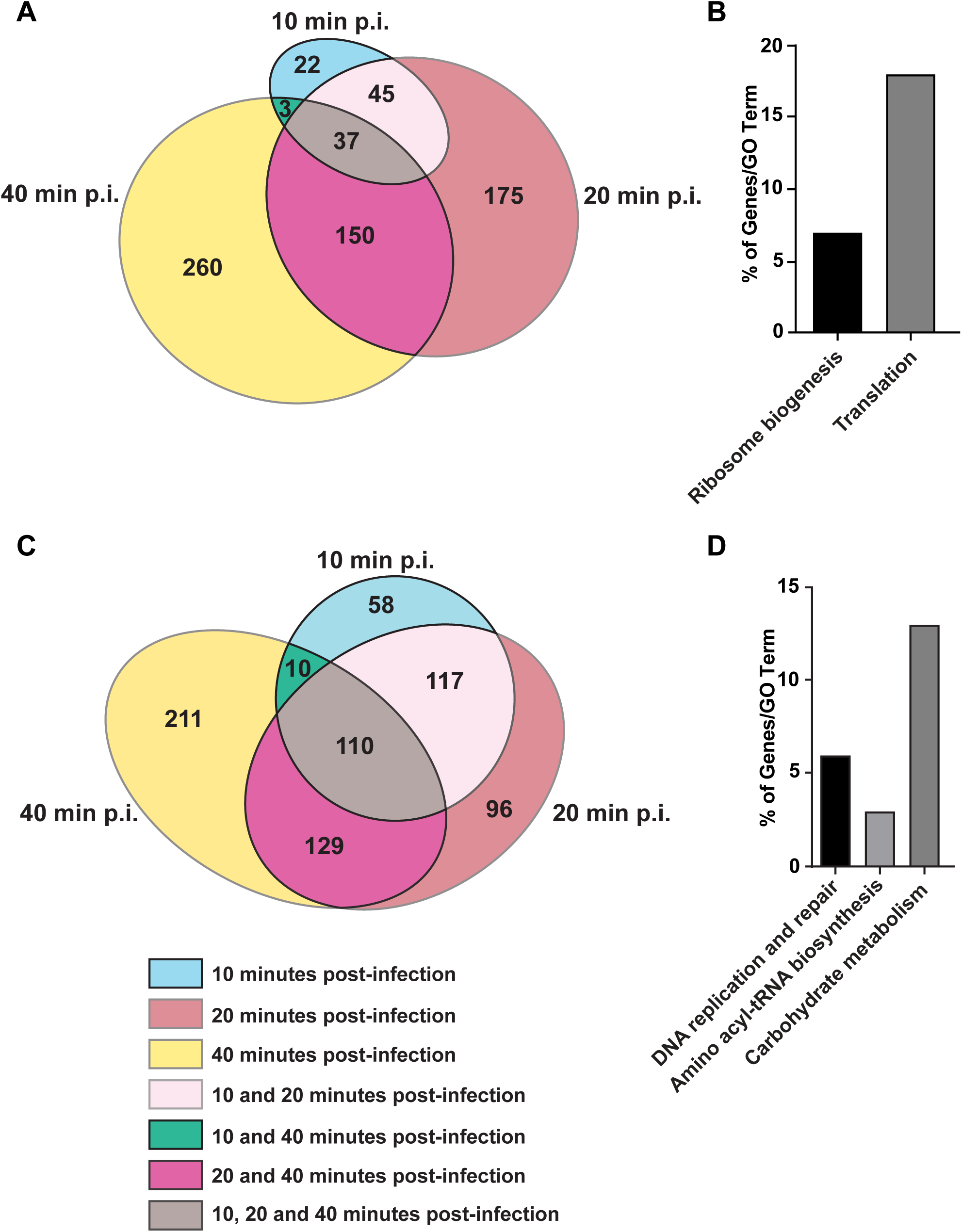
VPE25 modulation of *E. faecalis* genes during infection. **(A)** Euler diagram representing the number of host transcripts with reduced abundance during phage infection. **(B)** Among the 37 genes that are downregulated throughout the entire phage infection cycle, a large percentage belong to ribosome biogenesis and translation. **(C)** Phage induced expression of multiple host transcripts include those involved in **(D)** DNA replication and repair, tRNA biosynthesis and carbohydrate metabolism.

**Figure S6.**
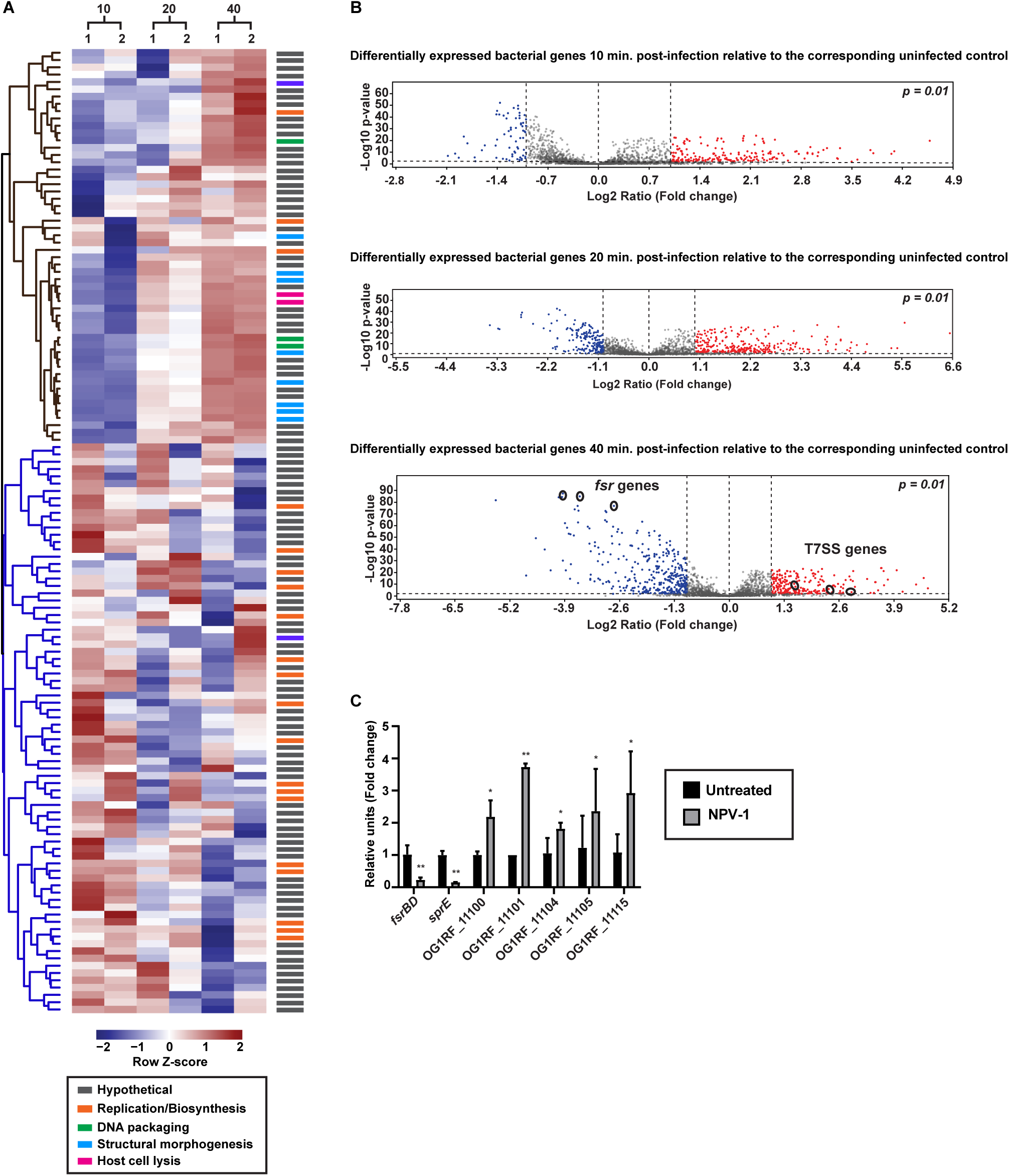
Transcriptomic profiling reveals alterations to phage and bacterial gene expression patterns throughout the course of infection. **(A)** Hierarchical clustering of differentially expressed VPE25 transcripts after 10, 20 and 40 minutes relative to 0 minutes post viral infection. 1 and 2 designate individual biological replicates. The transcripts are broadly classified into early and late gene clusters depicted in purple and green, respectively. **(B)** Volcano plots demonstrate changes in the gene expression pattern of *E. faecalis* OG1RF during VPE25 infection. Volcano plots highlight differentially expressed genes in the bacteria at 10 min (upper), 20 min (middle), and 40 min (bottom) post VPE25 infection. Downregulated genes are shown in blue and upregulated genes are in red (fold change > 2, *P* < 0.01). **(C)** Phage NPV1 infection downregulates *fsr* quorum sensing and upregulates T7SS in *E. faecalis* OG1RF. The Pip_EF_ independent phage NPV1 was used to query the expression of select *fsr* associated genes (*fsrBD* and *sprE*) and type VII secretion genes (*OG1RF_11100*, *OG1RF_11101*, *OG1RF_11104*, *OG1RF_11105* and *OG1RF_11115*) by qPCR 1 hour after NPV1 infection of *E. faecalis* OG1RF. The data are expressed as fold change of normalized mRNA in comparison to the uninfected controls and represent an average of three biological replicates ± the standard deviation. \**P* < 0.0001 and **P < 0.00001 by unpaired Student’s t-test.

Table S1. Differentially abundant transposon mutants during VPE25 selection of the *E. faecalis* OG1RF pooled Tn library.

Table S2. (A) Differential expression ratio of VPE25 genes during E. faecalis OG1RF infection relative to the start of infection. (B) Differential expression ratio of E. faecalis OG1RF genes during VPE25 infection relative to the untreated cultures.

Table S3. (A) GO and KEGG annotations of enterococcal genes downregulated throughout the course of VPE25 infection. (B) GO and KEGG annotations of enterococcal genes upregulated during VPE25 infection.

Table S4. Bacterial strains, phages, plasmids and primers used in this study.

